# A Benchmark for Virus Infection Reporter Virtual Staining in Fluorescence and Brightfield Microscopy

**DOI:** 10.1101/2024.08.30.610499

**Authors:** Maria Wyrzykowska, Gabriel della Maggiora, Nikita Deshpande, Ashkan Mokarian, Artur Yakimovich

## Abstract

Detecting virus-infected cells in light microscopy requires a reporter signal commonly achieved by immunohistochemistry or genetic engineering. While classification-based machine learning approaches to the detection of virus-infected cells have been proposed, their results lack the nuance of a continuous signal. Such a signal can be achieved by virtual staining. Yet, while this technique has been rapidly growing in importance, the virtual staining of virus-infected cells remains largely uncharted. In this work, we propose a benchmark and datasets to address this. We collate microscopy datasets, containing a panel of viruses of diverse biology and reporters obtained with a variety of magnifications and imaging modalities. Next, we explore the virus infection reporter virtual staining (VIRVS) task employing U-Net and pix2pix architectures as prototypical regressive and generative models. Together our work provides a comprehensive benchmark for VIRVS, as well as defines a new challenge at the interface of Data Science and Virology.

## Introduction

Animal and human cells and tissues appear achromatic under a conventional brightfield light microscope, making individual components hard to distinguish. Staining of such samples using conventional or fluorescent dyes serves an important role in understanding biochemical mechanisms, disease diagnosis, or tracking pathogen transmission (1, 2). Not only does staining improve the contrast for microscopic examination, but functional approaches like fluorescent immunohistochemistry allow for selective highlighting of the meaningful molecules in the sample (3, 4). Some staining techniques require treating samples with various buffers, reagents and dyes that selectively bind to cellular components like lipids, proteins, carbohydrates or nucleic acids (4, 5). Others employ genetic engineering to incorporate fluorescent proteins into the specimen (6). However, such sample preparation is complex and demands significant time and resources, as well as expertise and can potentially harm delicate samples or bias the results (7). To address this, the scientific community has explored virtual staining algorithms rooted in deep neural networks (DNN) that produce labelled images from label-free microscopy (8, 9). A multitude of histological stain types has been successfully replicated (8), often leveraging Convolutional Neural Networks (CNNs) as a staple technique in Computer Vision. Unlike classification, functional staining of proteins of interest akin to fluorescence microscopy allows visualising more subtle phenotypic nuances in biological images, including the shape and intensity of the signal. These features bear crucial biomedical information, including severity and multiplicity of infection.

Virtual staining can be formulated as either a regression or generation problem. Regression tasks produce deterministic outputs minimising the pixel-wise error, while generative models aim to learn the data distribution producing stochastic outputs (10). This implies that, while regression allows for more precise control over the pixel-to-pixel conversion, generative models can produce more diverse and realistic images. A popular choice for solving regression tasks is the U-Net (11) architecture based on the encoder-decoder structure (8, 9, 12, 13). For the image generation task in virtual staining, the generative adversarial networks (GANs) have been dominant (8, 9, 12, 13). In a GAN architecture (14), one network called the generator learns to generate the desired output images, while a second network — the discriminator — learns to distinguish between the GT images and the images produced by the generator. Examples include virtual staining of autopsy tissue (12), as well as multi-staining (13)

While both regression and generation-based virtual staining have been successfully applied to Digital Pathology (8, 9, 13) and microscopy (15, 16), a systematic benchmark for virtual staining of virus-infected cells has thus far not been established. At the same time, the classification of cells infected by nucleotropic viruses like human adenovirus (HAdV) and herpes simplex virus (HSV) based on an unspecific nuclear stain has been previously demonstrated (17, 18). This suggests that the morphological inductive biases relevant to virus infection of cells are present in the images of infected and uninfected cell nuclei. Image classification alone does not provide a similar amount of nuance as virtual staining. For example, infected cell morphology and the intensity of the fluorescent reporter expressed during virus infection correlate with the duration and severity of infection (19). Building models and benchmarks for virtual infection staining can significantly advance single-cell biology and aid in pathogen research or differential diagnostics.

To address this, we propose the Virus Infection Reporter Virtual Staining (VIRVS) benchmark consisting of infected cell microscopy datasets and metrics on typical models for regression and generation tasks. The datasets consist of high-content multi-channel fluorescence and brightfield microscopy of 4 different cell types infected by 5 different viruses spanning 2 different levels of magnification. Specifically, the VIRVS benchmark includes Human Adenovirus (HAdV), Herpes simplex virus 1 (HSV), vaccinia virus (VACV), influenza A virus (IAV) and human rhinovirus (RV) (Fig. 1). HAdV is a non-enveloped virus with an icosahedral capsid replicating exclusively in the nucleus (20). HSV is another example of a nucleotropic virus (21) we added to VIRVS. However, unlike HAdV, HSV is enveloped. VACV, IAV and RV replicate in the cytoplasm. VACV is a prototypic member of poxviruses – a family of large, enveloped double-stranded DNA viruses (22, 23). Together, the panel of the viruses used in VIRVS spans a variety of viruses with various tropisms, biology and lifecycles.

**Fig. 1.**
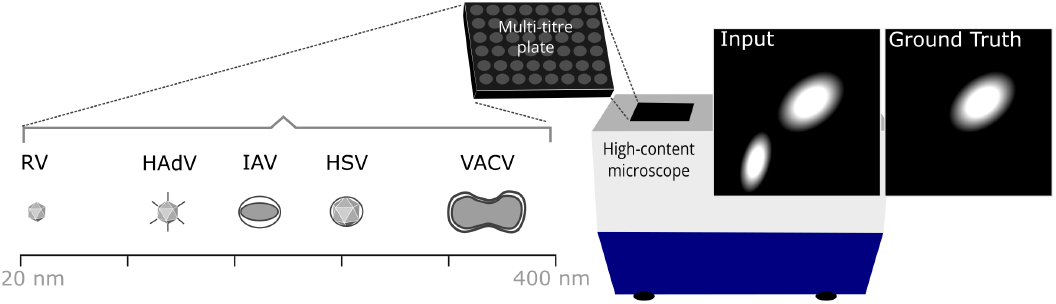
Schematic depiction of dataset sources and composition including Human Adenovirus (HAdV), Herpes simplex virus 1 (HSV), vaccinia virus (VACV), influenza A virus (IAV) and human rhinovirus (RV). HAdV is non-eveloped, approximately 90 nm in diameter and has a double-stranded DNA genome of around 36 kbp, replicates in the nucleus (28, 29). HSV is enveloped with an icosahedral capsid, 155 to 240 nm in diameter (30), has a double-stranded linear DNA genome that is approximately 152 kbp in length (21). VACV is enveloped, its genome is approximately 190 kbp and the dimensions of the brick-shaped capsid-containing virions are roughly 360 × 270 × 250 nm (31). IAV is enveloped, with an elliptical-shaped capsid of approximately 80 120 nm in diameter (32), has a segmented negative-sense RNA genome and replicates in the cytoplasm (33). RV is non-enveloped, 30-40 nm in size, replicates in the cell cytoplasm and has an icosahedral capsid, as well as a single-stranded positive-sense RNA genome of 7.2 to 8.5 kb in length (34, 35).

The multi-channel images used in the VIRVS benchmark are well registered and contain specific virus infection fluorescence signal used as ground truth (GT, *Y*) and unspecific fluorescence or brightfield signals used as input (*X*). The microscopy images were collated using the data from published works (24–26). The aim of the virtual staining models is to reproduce the GT as prediction Ŷ, being as close as possible to *Y*. To demonstrate the performance of regression models we employed a widely-used U-Net architecture (11). As a representative generative model, we employed the pix2pix (27). We evaluated whether regression or generation are the most adequate approaches in addressing this task, performing experiments on the aforementioned datasets spanning a broad range of characteristics and pathogens.

## Results

### VIRVS Dataset Collation

To establish the VIRVS benchmark we have collated datasets containing multi-channel high-content microscopy of virus-infected cells (24–26). These datasets include virus-specific reporter channel, for example green fluorescence protein expressed by the virus infected cells, and an unspecific channel allowing to visualise all cells in the field of view. In some datasets the unspecific channel includes nuclear staining, in other cases brightfield microscopy. Virus-specific reporter channel was reserved as GT, while unspecific was used as input (see Fig. 1).

Specifically, to devise a dataset for HAdV we employed data from (24). In this dataset the nuclei channel represents Hoechst 33342 staining and the viral channel indicates GFP concentration. Additionally, brightfield microscopy images of the cell monolayer were available. Infection was performed in the A549 human lung carcinoma-derived cell line. In the case of VACV we obtained the data from (25), where GFP expression from a transgenic VACV indicated infection in the monkey kidney BSC40 cell line, while the brightfield modality allowed to visualise all the cells. As several adjacent fields of view were acquired per well, we performed an overlap-based image stitching ensuring broader resampling. HAdV and VACV datasets were obtained at 10x and 4x magnification respectively. In both cases microscopy was performed in a live-cell imaging mode, where images were obtained as a time-lapse series, visualising the progression of infection across multiple time points. To avoid bias we have selected timepoints 49 (87 hours post infection) for HAdV and 100 (50h), 108 (54h) and 115 (57.5h) for VACV.

Data for the IAV, RV and HSV was obtained from a small molecule screening dataset performed at 4x magnification (26). HeLa Ohio Geneva cell line was used for RV, and A549 for IAV and HSV. Unlike HAdV and VACV, these datasets were obtained from fixed end-point assays. To avoid introducing morphological variations from the small molecule perturbations, we used exclusively control measurements containing virus-infected cells under mock conditions (dimethyl sulfoxide treatment). All images were obtained at 4x magnification and contained nuclear and virus channels. The source of virus signal in RV and HSV was GFP, while in the case of IAV the signal was obtained immunohistochemically using an anti-nucleoprotein (*α*-NP) antibody (see (26) for further details).

Notably, while HAdV and HSV are nucleotropic, affecting the nuclei phenotype visibly in microscopy images, VACV, IAV, and RV replicate in the cell cytoplasm. Given that only a nuclear channel is available as input for IAV and RV we presumed the inductive bias to be weaker or absent. Together, these datasets represent a wide range of imaging and infection characteristics, constituting a comprehensive benchmark of the models’ capabilities. Detailed descriptions of the number of images and resolution are presented in Table 1. For each dataset, we employed a data processing pipeline, resulting in training, validation, and test splits (see Methods section).

**Table 1.**
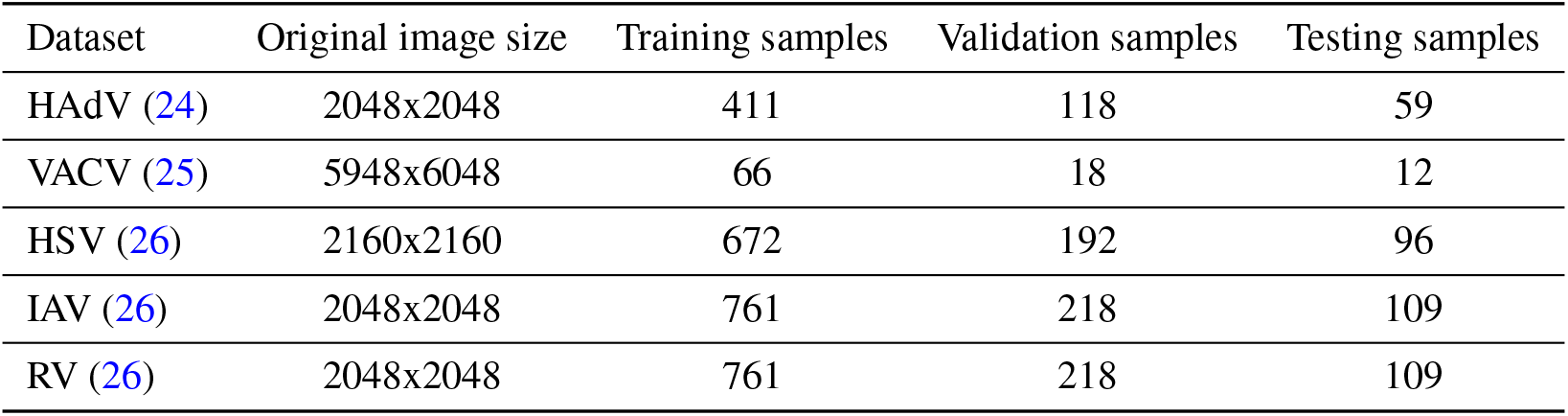
Final sizes of the datasets, including size of an individual image and number of images in training, validation and testing set for HAdV, VACV, HSV, IAV and RV data.

| Dataset | Original image size | Training samples | Validation samples | Testing samples |
| --- | --- | --- | --- | --- |
| HAdV (24) | 2048x2048 | 411 | 118 | 59 |
| VACV (25) | 5948x6048 | 66 | 18 | 12 |
| HSV (26) | 2160x2160 | 672 | 192 | 96 |
| IAV (26) | 2048x2048 | 761 | 218 | 109 |
| RV (26) | 2048x2048 | 761 | 218 | 109 |

### VIRVS Benchmark for Human Adenovirus

To evaluate the performance of the state-of-the-art virtual staining architectures on these data, we trained and evaluated U-Net (11) and pix2pix (27) models. For HAdV, we initially trained models on fluorescence microscopy images of nuclei. Subsequently, we trained separate models on combined fluorescence and brightfield images (HAdV (2ch)) to investigate the inductive bias introduced by cell shape visible only there. This approach was motivated by the observation that virus infection causes a cytopathic effect (CPE) a visible change in cell morphology (36–38). Furthermore, given that the GFP signal in all the datasets is localised in the cytoplasm, the quality of the resulting staining images is likely to be influenced by the cell shape. Additionally, we trained models solely on transmission light images, but the results were significantly worse, indicating that nuclear morphology is important for identifying the HAdV infection. This aligns with the understanding that HAdV, being a nucleotropic virus, affects the morphology of infected nuclei.

Fig. 2 illustrates examples of predictions made by the pix2pix and U-Net models for the HAdV data. Consistent with previous work, the nuclear signal of infected cells appears visually distinct (Fig. 2a, zoomed-in insets) suggesting a presence of a strong inductive bias. Model prediction results suggested that while both architectures commit similar errors, the pix2pix predictions are visually closer to the GT, capturing texture, shapes, and detail with greater fidelity.

**Fig. 2.**
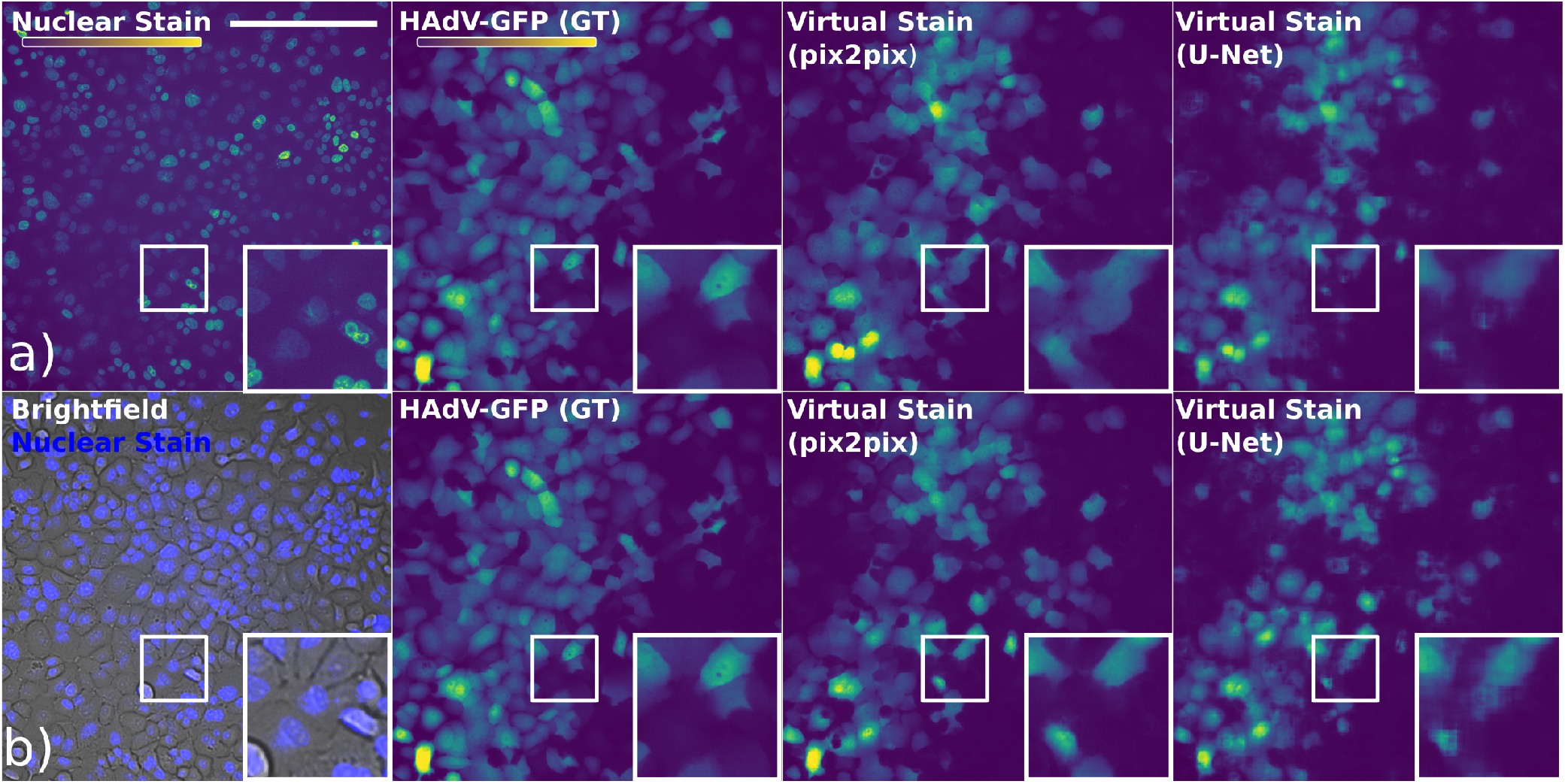
Examples of predictions made by the pix2pix and U-Net models for the HAdV data. The top row (a) displays the nuclear channel, GT and the virtual stains predicted by pix2pix and U-Net trained only on the nuclear channel. The bottom row (b) shows input with the nuclear channel superimposed on the brightfield channel (HAdV (2ch)), GT and the corresponding virtual stains predicted by the models trained on both channels. The scale bar is 500 *μ*m. Insets show respective zoomed-in regions. The colour code depicted in the colour bar.

To evaluate the model performance quantitatively, we computed image-wide metrics commonly used in image-to-image tasks, including mean square error (MSE), peak signal-to-noise ratio (PSNR) (39), and structural similarity index (SSIM) (40). To understand the models’ behaviour in a more comprehensive way, we additionally employed a rule-based approach to classify cell nuclei as either infected or healthy (see Methods section). We report classification and segmentation metrics such as IoU, F1, accuracy, precision and recall based on the results, summarised in the Table 2. Furthermore, in the Supplement, we report additional results: MSE, PSNR and SSIM calculated separately for the background and fore-ground sections of the images in Table 4, IoU, F1, accuracy, precision and recall calculated for whole cells for HAdV data in Table 5 and means and standard deviations calculated for every metric across the data points in Table 6 and 7.

**Table 2.** Performance of U-Net and pix2pix models, trained on HAdV, HAdV (2ch), VACV, HSV, IAV and RV data. Metrics that we report are MSE, PSNR, SSIM, IoU, F1, accuracy, precision and recall. For pix2pix, we report the mean and standard deviation of each metric based on 3 runs.

| Dataset | Model | MSE (↓) | PSNR (↑) | SSIM (↑) | IoU (↑) | F1 (↑) | Accuracy (↑) | Precision (↑) | Recall (↑) |
| --- | --- | --- | --- | --- | --- | --- | --- | --- | --- |
| HAdV | pix2pix | $0.034 \pm 5.0\text{e-}05$ | $24.214 \pm 7\text{e-}03$ | $0.870 \pm 3.0\text{e-}05$ | $0.300 \pm 7.0\text{e-}04$ | $0.437 \pm 6.0\text{e-}04$ | $0.959 \pm 4.0\text{e-}05$ | $0.503 \pm 7.0\text{e-}04$ | $0.473 \pm 1.0\text{e-}03$ |
|  | U-Net | **0.031** | **25.331** | **0.899** | 0.268 | 0.404 | **0.963** | **0.557** | 0.357 |
| HAdV (2ch) | pix2pix | **$0.032 \pm 4.0\text{e-}05$** | **$24.824 \pm 1.1\text{e-}02$** | $0.892 \pm 1.0\text{e-}04$ | **$0.239 \pm 8.0\text{e-}04$** | **$0.364 \pm 1.0\text{e-}03$** | $0.960 \pm 8.0\text{e-}05$ | **$0.524 \pm 1.0\text{e-}03$** | **$0.322 \pm 2.0\text{e-}03$** |
|  | U-Net | **0.032** | 24.767 | **0.895** | 0.235 | 0.360 | **0.961** | 0.522 | 0.306 |
| VACV | pix2pix | $0.083 \pm 4.0\text{e-}05$ | $21.087 \pm 1.5\text{e-}02$ | $0.825 \pm 5.0\text{e-}05$ | - | - | - | - | - |
|  | U-Net | **0.057** | **24.563** | **0.860** | - | - | - | - | - |
| HSV | pix2pix | $0.016 \pm 9.0\text{e-}07$ | $27.514 \pm 2.0\text{e-}03$ | $0.917 \pm 1.0\text{e-}05$ | $0.660 \pm 2.0\text{e-}04$ | $0.794 \pm 1.0\text{e-}04$ | $0.985 \pm 6.0\text{e-}06$ | **$0.862 \pm 3.0\text{e-}05$** | $0.741 \pm 3.0\text{e-}04$ |
|  | U-Net | **0.010** | **29.673** | **0.938** | **0.727** | **0.841** | **0.988** | 0.858 | **0.830** |
| IAV | pix2pix | $0.054 \pm 1.0\text{e-}05$ | $22.266 \pm 4.0\text{e-}03$ | $0.875 \pm 2.0\text{e-}05$ | **$0.249 \pm 1.0\text{e-}04$** | **$0.391 \pm 9.0\text{e-}05$** | $0.967 \pm 2.0\text{e-}05$ | $0.485 \pm 3.0\text{e-}04$ | **$0.341 \pm 3.0\text{e-}04$** |
|  | U-Net | **0.041** | **24.374** | **0.905** | 0.174 | 0.290 | **0.971** | **0.655** | 0.190 |
| RV | pix2pix | $0.045 \pm 6.0\text{e-}06$ | $21.668 \pm 1.0\text{e-}04$ | $0.863 \pm 1.0\text{e-}05$ | $0.219 \pm 2.0\text{e-}04$ | $0.357 \pm 3.0\text{e-}04$ | $0.963 \pm 2.0\text{e-}05$ | $0.347 \pm 4.0\text{e-}04$ | **$0.379 \pm 3.0\text{e-}04$** |
|  | U-Net | **0.024** | **24.738** | **0.917** | **0.294** | **0.448** | **0.978** | **0.614** | 0.358 |

**Table 3.** Percentile values used to normalize the ground truth (GT), nuclear stain and brightfield channels for HAdV, VACV, HSV, IAV and RV datasets.

| Dataset | GT | GT | Nuclear stain | Nuclear stain | Brightfield | Brightfield |
| --- | --- | --- | --- | --- | --- | --- |
| HAdV | 0.0104 | 0.0741 | 0.0035 | 0.0178 | 0.032 | 0.4089 |
| VACV | 0.0067 | 1.0 | - | - | 0.4956 | 1.0 |
| HSV | 0.0033 | 0.1926 | 0.0074 | 0.2907 | - | - |
| IAV | 0.0091 | 0.1769 | 0.011 | 0.2156 | - | - |
| RV | 0.0082 | 0.1712 | 0.0086 | 0.2193 | - | - |

**Table 4.**
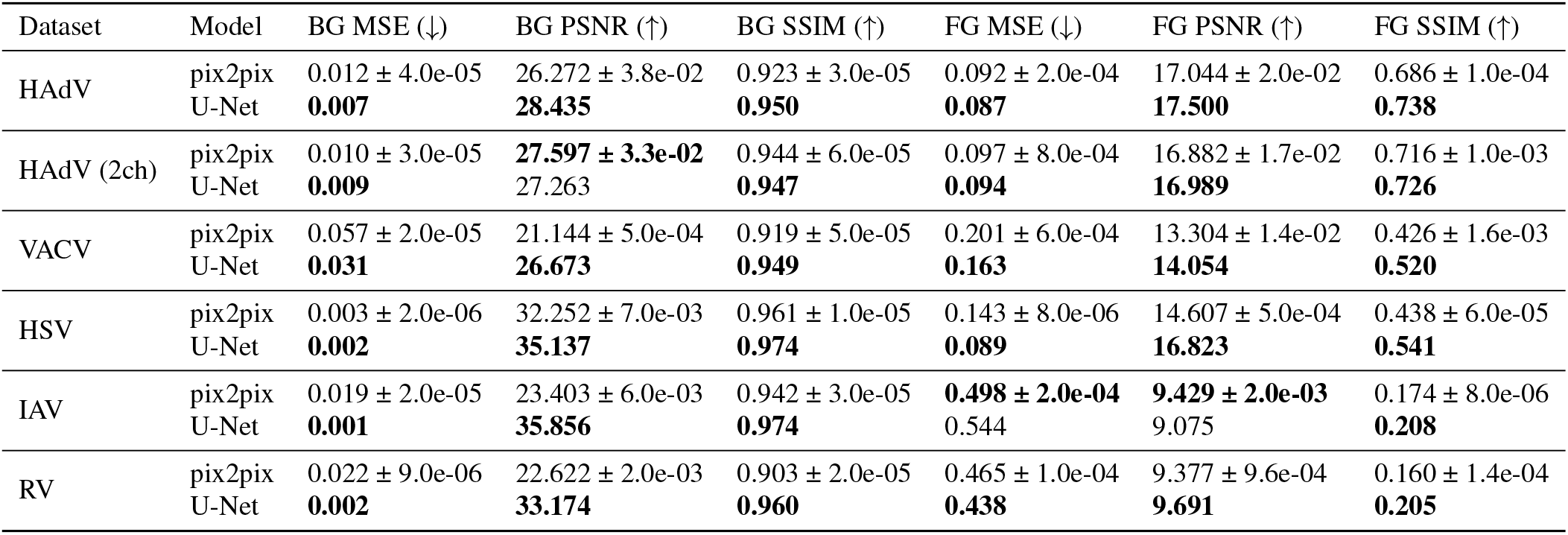
Performance of U-Net and pix2pix models, trained on HAdV, HAdV (2ch), VACV, HSV, IAV and RV data. Metrics reported are MSE, PSNR and SSIM, calculated separately for background (BG) or foreground (FG). For pix2pix, we report the mean and standard deviation of each metric based on the 3 trials.

**Table 5.** Performance of U-Net and pix2pix models, trained on HAdV and HAdV (2ch) data. Metrics reported are IoU, F1, accuracy, precision, and recall, calculated based on cells segmentation masks. For pix2pix, we report the mean and standard deviation of each metric based on 3 trials.

| Dataset | Model | IoU (↑) | F1 (↑) | Accuracy (↑) | Precision (↑) | Recall (↑) |
| --- | --- | --- | --- | --- | --- | --- |
| HAdV | pix2pix | **$0.326 \pm 6.0\text{e-}04$** | **$0.462 \pm 6.0\text{e-}04$** | $0.856 \pm 2.0\text{e-}04$ | $0.500 \pm 7.0\text{e-}04$ | **$0.548 \pm 6.0\text{e-}04$** |
|  | U-Net | 0.303 | 0.440 | **0.871** | **0.528** | 0.447 |
| HAdV (2ch) | pix2pix | **$0.294 \pm 2.0\text{e-}03$** | **$0.428 \pm 2.0\text{e-}03$** | $0.868 \pm 4.0\text{e-}04$ | **$0.529 \pm 3.0\text{e-}03$** | **$0.438 \pm 2.0\text{e-}03$** |
|  | U-Net | 0.282 | 0.413 | **0.871** | 0.516 | 0.408 |

**Table 6.** Performance of U-Net and pix2pix models, trained on HAdV, HAdV (2ch), VACV, HSV, IAV, and RV data. Metrics reported are the mean and standard deviation between results for each datapoint for MSE, PSNR, and SSIM, along with their foreground (FG) and background (BG) versions.

| Dataset | Model | MSE (↓) | PSNR (↑) | SSIM (↑) | BG MSE (↓) | BG PSNR (↑) | BG SSIM (↑) | FG MSE (↓) | FG PSNR (↑) | FG SSIM (↑) |
| --- | --- | --- | --- | --- | --- | --- | --- | --- | --- | --- |
| HAdV | pix2pix | 0.034 ± 0.032 | 24.214 ± 4.218 | 0.870 ± 0.060 | 0.012 ± 0.009 | 26.272 ± 3.267 | 0.923 ± 0.016 | 0.092 ± 0.055 | 17.044 ± 2.433 | 0.686 ± 0.084 |
|  | U-Net | **0.031 ± 0.038** | **25.331 ± 4.753** | **0.899 ± 0.067** | **0.007 ± 0.006** | **28.435 ± 3.112** | **0.950 ± 0.011** | **0.087 ± 0.060** | **17.500 ± 2.796** | **0.738 ± 0.076** |
| HAdV (2ch) | pix2pix | **0.032 ± 0.032** | **24.824 ± 4.776** | 0.892 ± 0.065 | 0.010 ± 0.007 | **27.597 ± 3.954** | 0.944 ± 0.012 | 0.097 ± 0.057 | 16.882 ± 2.698 | 0.716 ± 0.083 |
|  | U-Net | **0.032 ± 0.034** | 24.767 ± 4.485 | **0.895 ± 0.067** | **0.009 ± 0.006** | 27.263 ± 3.056 | **0.947 ± 0.009** | **0.094 ± 0.057** | **16.989 ± 2.542** | **0.726 ± 0.078** |
| VACV | pix2pix | 0.083 ± 0.070 | 21.087 ± 3.593 | 0.825 ± 0.096 | 0.057 ± 0.069 | 21.144 ± 4.480 | 0.919 ± 0.073 | 0.201 ± 0.078 | 13.304 ± 1.621 | 0.426 ± 0.049 |
|  | U-Net | **0.057 ± 0.047** | **24.563 ± 7.304** | **0.860 ± 0.090** | **0.031 ± 0.048** | **26.673 ± 6.911** | **0.949 ± 0.059** | **0.163 ± 0.044** | **14.054 ± 1.195** | **0.520 ± 0.068** |
| HSV | pix2pix | 0.016 ± 0.007 | 27.514 ± 2.647 | 0.917 ± 0.030 | 0.003 ± 0.002 | 32.252 ± 2.108 | 0.961 ± 0.015 | 0.143 ± 0.037 | 14.607 ± 1.055 | 0.438 ± 0.062 |
|  | U-Net | **0.010 ± 0.005** | **29.673 ± 2.714** | **0.938 ± 0.023** | **0.002 ± 0.002** | **35.137 ± 2.704** | **0.974 ± 0.010** | **0.089 ± 0.034** | **16.823 ± 1.503** | **0.541 ± 0.073** |
| IAV | pix2pix | 0.054 ± 0.014 | 22.266 ± 1.103 | 0.875 ± 0.023 | 0.019 ± 0.004 | 23.403 ± 0.855 | 0.942 ± 0.007 | **0.498 ± 0.211** | **9.429 ± 1.818** | 0.174 ± 0.088 |
|  | U-Net | **0.041 ± 0.016** | **24.374 ± 1.861** | **0.905 ± 0.023** | **0.001 ± 0.000** | **35.856 ± 1.587** | **0.974 ± 0.003** | 0.544 ± 0.235 | 9.075 ± 1.887 | **0.208 ± 0.109** |
| RV | pix2pix | 0.045 ± 0.009 | 21.668 ± 0.927 | 0.863 ± 0.019 | 0.022 ± 0.004 | 22.622 ± 0.818 | 0.903 ± 0.011 | 0.465 ± 0.056 | 9.377 ± 0.546 | 0.160 ± 0.028 |
|  | U-Net | **0.024 ± 0.007** | **24.738 ± 1.400** | **0.917 ± 0.018** | **0.002 ± 0.001** | **33.174 ± 1.650** | **0.960 ± 0.007** | **0.438 ± 0.088** | **9.691 ± 0.888** | **0.205 ± 0.053** |

**Table 7.** Performance of U-Net and pix2pix models, trained on HAdV, HAdV (2ch), HSV, IAV, and RV data. Metrics reported are the mean and standard deviation between results for each datapoint for IoU, F1, accuracy, precision, and recall calculated based on the nuclei masks.

| Dataset | Model | IoU (↑) | F1 (↑) | Accuracy (↑) | Precision (↑) | Recall (↑) |
| --- | --- | --- | --- | --- | --- | --- |
| HAdV | pix2pix | **0.300 ± 0.163** | **0.437 ± 0.199** | 0.959 ± 0.022 | 0.503 ± 0.302 | **0.473 ± 0.141** |
|  | U-Net | 0.268 ± 0.137 | 0.404 ± 0.174 | 0.963 ± 0.025 | **0.557 ± 0.309** | 0.357 ± 0.124 |
| HAdV (2ch) | pix2pix | **0.239 ± 0.138** | **0.364 ± 0.182** | 0.960 ± 0.025 | **0.524 ± 0.320** | **0.322 ± 0.141** |
|  | U-Net | 0.235 ± 0.140 | 0.360 ± 0.183 | **0.961 ± 0.026** | 0.522 ± 0.321 | 0.306 ± 0.129 |
| HSV | pix2pix | 0.660 ± 0.054 | 0.794 ± 0.040 | 0.985 ± 0.005 | **0.862 ± 0.052** | 0.741 ± 0.063 |
|  | U-Net | **0.727 ± 0.048** | **0.841 ± 0.034** | **0.988 ± 0.005** | 0.858 ± 0.050 | **0.830 ± 0.056** |
| IAV | pix2pix | **0.249 ± 0.082** | **0.391 ± 0.107** | 0.967 ± 0.008 | 0.485 ± 0.154 | **0.341 ± 0.095** |
|  | U-Net | 0.174 ± 0.067 | 0.290 ± 0.098 | **0.971 ± 0.010** | **0.655 ± 0.186** | 0.190 ± 0.072 |
| RV | pix2pix | 0.219 ± 0.047 | 0.357 ± 0.062 | 0.963 ± 0.015 | 0.347 ± 0.088 | **0.379 ± 0.056** |
|  | U-Net | **0.294 ± 0.085** | **0.448 ± 0.104** | **0.978 ± 0.008** | **0.614 ± 0.083** | 0.358 ± 0.102 |

In our analysis, the U-Net model showed superior results for the HAdV dataset when trained on a single nuclear channel, with a lower MSE, and a higher PSNR, SSIM, accuracy and precision compared to pix2pix. However, for the combination of brightfield and nuclear channels (HAdV (2ch)), pix2pix slightly outperformed U-Net in most metrics, despite having a marginally lower SSIM and accuracy. These findings suggest that pix2pix tends to overestimate the GFP signal, leading to higher recall but worse or comparable precision, while U-Net delivers better accuracy, especially when excluding the brightfield channel.

Together with the visual examination, the results suggest that while pix2pix provided more realistic-looking cell shapes, this did not significantly improve the metrics such as MSE, PSNR or SSIM of the models for HAdV, which remained close to each other. The difference is more noticeable when metrics such as IoU, precision and recall are incorporated. Notably, the CPE in the brightfield images was not very pronounced at this stage of infection, which might have limited the inductive bias and introduced unnecessary noise, disrupting U-Net’s performance.

### VIRVS Benchmark for Vaccinia Virus

To further explore if CPE in brightfield microscopy can provide a robust inductive bias for virtual staining we evaluated the performance of U-Net and pix2pix in predicting virus signal of VACV, known to induce a strong CPE (41). Noteworthy, the CPE is visible prior to the GFP signal, as it takes time to express and accumulate GFP molecules in the infected cell. Fig. 3 illustrates examples of predictions made by the pix2pix and U-Net for the VACV data. Both perform well on this dataset, producing signal comparable to the GT. U-Net underestimated both the magnitude and the extent of the signal (Fig. 3, dashed lines in the zoomed-in insets) compared to GT, while pix2pix overestimated the magnitude, overestimating the extent only slightly. Unsurprisingly, the predictions of both architectures suffered from the artefacts introduced by the stitching procedure, however, since they occurred in predictions of both architectures to a comparable degree, we assumed they did not significantly impact our analysis.

**Fig. 3.**
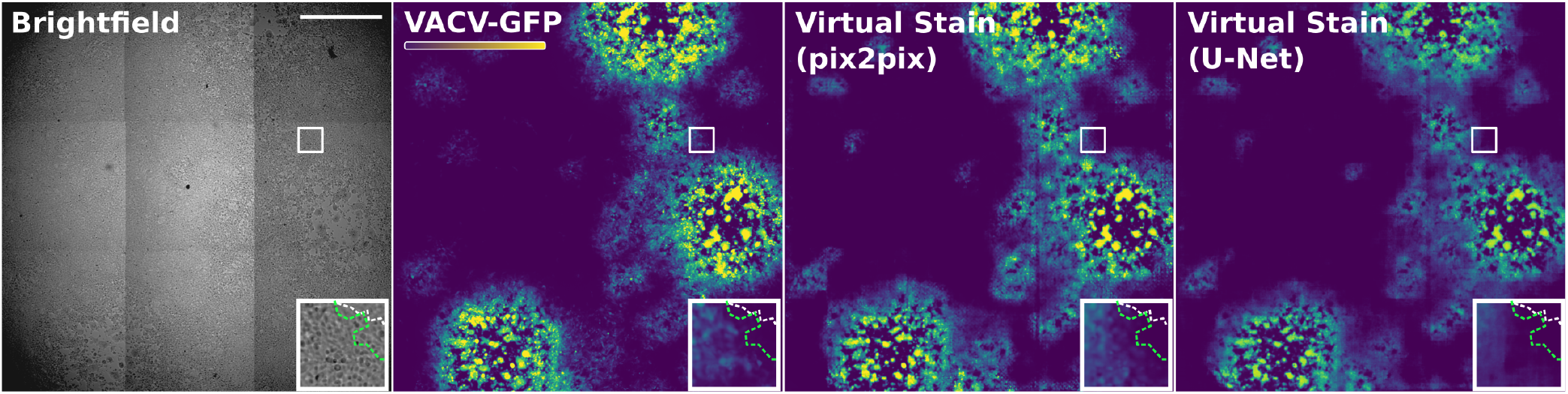
Examples of predictions made by the pix2pix and U-Net models for the VACV data. Starting from the left, the brightfield channel and GT are visualised, followed by pix2pix and U-Net predictions. Insets show respective zoomed-in regions. The dashed green line indicates the approximate edge of the GFP signal. Dashed white – an approximate edge of the cytopathic effect. The colour code is depicted in the colour bar. Scale bar 2000 *μ*m.

Looking at the metrics (Table 2), the U-Net model outperformed pix2pix. The U-Net achieved a lower MSE, higher PSNR, and higher SSIM, indicating superior overall performance. This may be attributed to the overestimation of the signal magnitude committed by the pix2pix. We do not report other metrics due to the lack of the fluorescence nuclei signal for VACV hampering the accurate segmentation in the dense cell monolayer.

### VIRVS Benchmark for HSV, IAV and RV

Finally, to explore potential inductive bias in the nuclear signal of cells infected with cytoplasm-replicating IAV and RV compared to nucleotropic HSV, we trained U-Net and pix2pix on a dataset derived from a small molecule screen (26). Figure 4a-c illustrates examples of predictions for the HSV, IAV and RV respectively. Notably, the nuclear staining of HSV data displayed strong intensity irregularities corresponding to the GT GFP signal, confirming the presence of a strong inductive bias. Unsurprisingly, both U-Net and pix2pix performed equally well visually on this task.

**Fig. 4.**
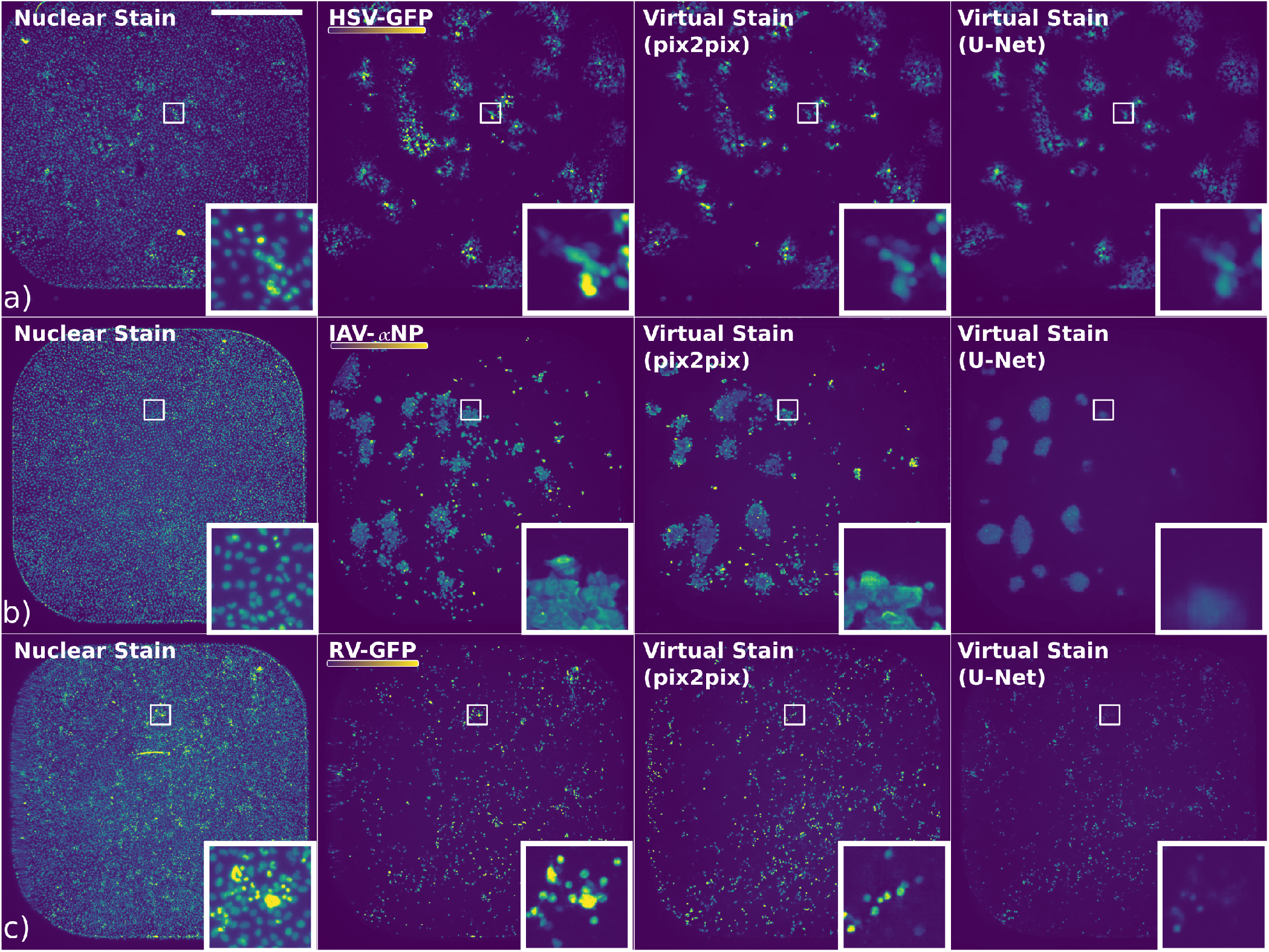
Examples of predictions made by the pix2pix and U-Net models for the HSV, IAV and RV data. In each row, starting from the left, the brightfield channel and GT are visualised, followed by pix2pix and U-Net predictions. The top row (a) displays the HSV data, the middle row (b) shows the IAV data, and the bottom row (c) presents the RV data. Scale bar 1000 *μ*m

Interestingly, IAV nuclear signal showed some irregularities corresponding to the virus signal as well, suggesting the presence of some inductive bias. As the intensity of the nuclear stain is directly connected to the cell cycle and given the role of IAV in host cell cycle arrest (42), it is tempting to speculate that the inductive bias facilitating the prediction may be the host DNA content. Curiously, while both pix2pix and U-Net demonstrated some visual overlap with the signal in the GT, their performance was weak. This manifested as blurry sporadic patches generated by the U-Net and multiple false positive pixels (hallucinations) generated by the pix2pix. At the same time, the performance of the models on RV proved to be consisting of mostly false positives. This suggested that in the absence of a strong inductive bias, both U-Net and pix2pix hallucinated the results.

In our evaluation of the HSV, IAV, and RV metrics (see Tab. 2), U-Net consistently outperformed pix2pix in terms of image prediction metrics. For the HSV dataset, U-Net achieved a better MSE, PSNR and SSIM, indicating superior similarity to GT images. It also achieved a high IoU, F1, accuracy and recall, with slightly worse precision than pix2pix. Analogously, in the IAV dataset, U-Net’s MSE, PSNR, SSIM, accuracy and precision surpassed pix2pix’s. On the other hand, pix2pix achieved significantly higher IoU, F1 and recall, suggesting that pix2pix may have overestimated the *α*NP signal in contrast to U-Net’s underestimation. For the RV dataset, U-Net continued to excel. Only recall was higher for pix2pix.

Consistent with our expectations, the nucleotropic HSV proved to be the easiest to predict based on the nuclear signal, reaching the best metrics overall. For U-Net, the performance of the RV was higher than the IAV according to all the metrics, except for the precision, contrarily to the visual impression, that suggested high hallucination rate, exhibiting that these metrics are vulnerable to hallucinations. RV may have outperformed IAV in these metrics due to the high false positive rate in RV predictions.

## Discussion

Detecting virus-infected cells in microscopy can help unveil novel host-pathogen interactions in the context of Cell Biology (24) and facilitate the discovery of novel antivirals in the context of high-content screening (25, 26). In this work, we demonstrate that the emerging field of deep-learning-based Virtual Staining can facilitate the detection of virus-infected cells from seemingly irrelevant signal. Compared to detection by classification (24), the ability to generate a pixel-level prediction provides a greater level of nuance including visual guidance to severity and duration of infection (see Fig. 3). Furthermore, we collate the datasets, baselines, and metrics into a benchmark VIRVS that is readily available for the community to download and improve upon.

Virtual Staining can be formulated as either regression or image generation. Using state-of-the-art architectures for these tasks, we demonstrate that in the case of VIRVS, both tasks are applicable provided sufficient inductive bias is present in the data. In our case, the generative model tended to yield more realistic-looking results, yet it often hallucinated where the signal wasn’t present in the GT. The regression model, in turn, always produced less realistic results with more conservative predictions. Overall, the stronger the inductive bias the better were the results of both generative and regression models.

From the metric perspective, the values of MSE, PSNR and SSIM suggest that overall U-Net demonstrated favourable performance across all viruses and image resolutions, except the HAdV (2ch) data, where pix2pix achieved comparable results. It is tempting to speculate, that this can be attributed to more conservative predictions of U-Net and a higher false-positive rate of pix2pix. Additional metrics, such as IoU, F1, accuracy, precision and recall demonstrated more nuance between the models and enabled more detailed comparison. Notably, performance on HSV images ranked highest overall, whereas VACV ranked lowest. Moreover, in contrast to the visual impression, pix2pix on VACV data performed the worst overall in all the metrics. Additionally, while pix2pix attained lower scores and sometimes hallucinated, its outputs appeared more realistic, detailed, and sharp compared to U-Net’s, which can be observed in HAdV or IAV predictions. This suggests that the commonly used metrics, such as MSE or PSNR may not always capture all the nuances of the virtual staining. The inclusion of metrics such as IoU, precision or recall could help alleviate this issue but computing these requires segmentation-level annotations. To thoroughly evaluate the model’s behaviour, one should consider all of these metrics in conjunction. This can be further enhanced by examining them at a finer level of detail, such as analysing them separately for the background and foreground of the image, as suggested in the Supplement. Additionally, a visual examination of the predictions should be incorporated to provide a more comprehensive understanding. Only then it is possible to gain a deep understanding of the performance.

While accuracy is of utmost importance for this task, overly cautious modelling or blurring out details to achieve the lowest error is insufficient. On the other hand, sacrificing truthfulness for more visually appealing predictions is unsound. Future works should focus on striking a balance between the advantages and shortcomings of the regressive and generative approach to this task and strive to find meaningful evaluation techniques to assess it. The VIRVS benchmark introduced in this work will be instrumental in these efforts. Through it, we demonstrated the feasibility of virtual staining for virus infection in cells for the first time. We hope that introducing these benchmark datasets together with our baseline characterisation will spur interest in exploring virtual staining in the context of viral infections.

## Code availability

Code has been published on GitHub.

## Data availability

Raw data used during the study can be found in corresponding references. Filtered and processed data have been published in RODARE (43).

## ACKNOWLEDGEMENTS

This work was partially funded by the Center for Advanced Systems Understanding (CASUS) which is financed by Germany’s Federal Ministry of Education and Research (BMBF) and by the Saxon Ministry for Science, Culture, and Tourism (SMWK) with tax funds on the basis of the budget approved by the Saxon State Parliament. MW was supported by the Scultetus Early Career Fellowship Program at CASUS.

The authors acknowledge the financial support by the Federal Ministry of Education and Research of Germany and by Sächsische Staatsministerium für Wissenschaft, Kultur und Tourismus in the programme Center of Excellence for AI-research „Center for Scalable Data Analytics and Artificial Intelligence Dresden/Leipzig”, project identification number: ScaDS.AI.

## Methods

### VACV and HAdV Data Processing

Each of VACV plaques was imaged to produce 9 files per channel, that need to be stitched to recreate the whole plaque. To achieve this, multiview-stitcher toolbox has been used. The stitching was first performed on the third channel, representing the brightfield microscopy image of the samples. Then, the parameters found for this channel were used to stitch the rest of the channels. VACV dataset represents a timelapse, from which timesteps 100, 108 and 115 have been selected to produce the data then used in the experiments. Images have been centercropped to 5948×6048 to match the size of the smallest image in the dataset (rounded down to the closest multiple of 2). The data was additionally manually filtered to remove the samples that constituted only uninfected cells (C02, C07, D02, D07, E02, E07, F02, F07).

HAdV dataset is also a timelapse, from which only the last timestep (49th) has been selected.

### HSV, IAV, RV Data Processing

For the rest of the datasets (HSV, IAV, RV) only the negative control data was used, which was selected in the following way: from the data collected at the University of Zürich, from the Screen samples only the first 2 columns were selected and from the ZPlates and prePlates samples only the first 12 columns.

All of the datasets were divided into training, validation and test holdouts in 0.7:0.2:0.1 ratios, using random seed 42 to ensure reproducibility. For the time-lapse data, it was ensured that the same sample from different timesteps only exists in one of the holdouts, to prevent information leakage and ensure fair evaluation. All of the samples were normalised to [-1, 1] range, by subtracting the 3rd percentile and dividing by the difference between percentile 99.8 and 3, clipping to [0, 1] and scaling to [-1, 1] range. For the brightfield channel of HAdV, percentiles 0.1 and 99.9 were used. These cutoff points were selected based on the analysis of the histograms of the values attained by the data, to make the best use of the available data range. Specific values used for the normalization are summarized in Figure 3.

### Model architecture

Pix2pix model was implemented as described in the original paper (27), based on official Tensorflow pix2pix tutorial, licensed under the Apache 2.0 License. The architecture of the U-Net model (44) was identical to the one used as the generator in pix2pix.

### Cellpose masks preparation

To prepare the cell nuclei masks, the Cellpose (45) model with pre-trained weights cyto3 (46) have been used on the fluorescence channel. The diameter was set to 7 for all the datasets except for HAdV, for which the automatic estimation of the diameter was employed. To prepare the cell nuclei masks, the Cellpose model with pre-trained weights cyto3 has been used on the fluorescence channel. Cell masks were prepared using Cellpose with pre-trained weights cyto3 with a diameter set to 70 on brightfield images stacked with fluorescence nuclei signal.

### Training setup

The size of the input expected by both of the models is 256×256 px. Random 256×256 crops of the data of the original size were used for training. For the model trained on both modalities of HAdV data, they were provided to the model as two separate channels. For both of the models, random jitter as suggested as an augmentation in pix2pix paper (27) was applied, in addition to random up and down and left and right flips with probability *p* = 0.5.

The hyperparameters used for training come from the pix2pix paper; Adam optimiser (47) with learning rate 2e-4 and *β*_1_ momentum parameter set to 0.5. The batch size was set to 8. The loss function used for pix2pix was the one suggested in the paper, using 0.5 as the weight of the discriminator loss. For UNet mean absolute error (MAE) was used.

All the models were trained for 100000 steps. A step is defined as a gradient descent update after processing one batch. A random seed equal to 42 was used to ensure reproducibility. All experiments were conducted on NVIDIA RTX A100 and V100 GPU devices.

### Evaluation setup

Models have been evaluated on the test set. For evaluation purposes, we used the whole images, center-cropped to the closest multiple of 256, as required by the model architecture. The batch size used for the evaluation is 1. Random seeds used during the evaluation were 42 for U-Net and 42, 43, 44 for pix2pix.

During evaluation, we used the batch normalization statistics of the test batch for both of the models. This approach, coined as Instance Normalization, was originally used in pix2pix, but also suggested for U-Net applied to biomedical data in (48). Additionally, for pix2pix the dropout layers are activated, as recommended in the pix2pix paper (27).

Metrics used to evaluate the performance are mean square error (MSE) (49), structural similarity index (SSIM) (40) and peak signal-to-noise ratio (PSNR) (50) and are implemented as follows:

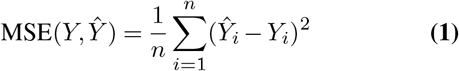

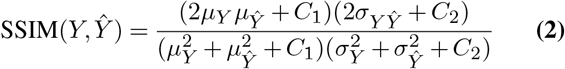

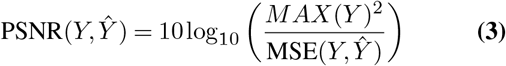

where *μ*_*Y*_ and *μ*_*Ŷ*_ are the average of *Y* and *Ŷ*, 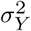 and 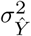 are the variances, *σ*_*YŶ*_ is the covariance of *Y* and *Ŷ*, and *C*_1_, *C*_2_ are constants as described in the original implementation of SSIM (40) and MAX_*Y*_ is the maximum possible data range in image, in our case 2.

The classification is performed through the following steps:

1. Cell nuclei are segmented using Cellpose.
2. For each segmented nucleus, we calculate the fraction of pixels in the GFP or *α*NP channel (either from ground truth or produced by the trained model) that have values higher than –0.9 for HAdV or –0.8 for other viruses. If this fraction exceeds 0.5, the nucleus is considered infected. These threshold values were selected based on a careful visual examination.
3. A mask is generated for each image. Background pixels (where no nucleus was segmented by Cellpose) and pixels of healthy nuclei are labelled with 0, while pixels of infected nuclei are labelled with 1.

These masks are created based on ground truth data, as well as predictions from the pix2pix and U-Net models. The overall accuracy is then calculated using the following formula:

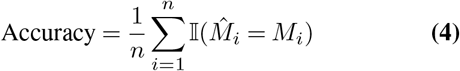

where *M* is the mask created based on ground truth data, 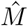 is the mask created based on the model’s prediction, *n* is the size of the image, and 𝕀(·) is the indicator function that equals 1 when its argument is true and 0 otherwise.

This, which we include for completeness sake, can be misleadingly high. This is because most of the pixels in the mask image typically do not correspond to any objects and are set to 0, regardless of the model’s prediction. For this reason, we exclude regions of the data without any objects from the calculation of the other metrics and accuracy should be interpreted alongside these other metrics.

Based on the masks, a confusion matrix for the binary classification of cell nuclei is created on a per-pixel basis, excluding pixels that do not belong to any cell nucleus. The number of true positives (TP), true negatives (TN), false positives (FP), and false negatives (FN) are then used to calculate the following metrics:

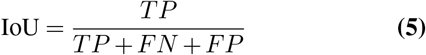

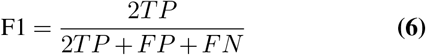

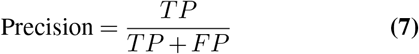

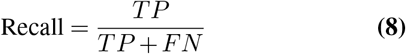

## Supplementary Note 1: Background-foreground metrics

To gain a more comprehensive understanding of the models’ behavior, we employed additional sets of metrics. We calculated traditional metrics on the data segmented into “foreground” and “background” using a threshold-based approach. Specifically, regions of an image with values higher than a certain manually chosen threshold in the ground truth viral signal channel were classified as “foreground,” while the rest were classified as “background.” The threshold was set to –0.8 for all viruses except HAdV, for which we used –0.9. Table 4 contains the results.

The results suggest that predicting the background is easier, as shown by lower MSE and higher PSNR and SSIM scores across all models. VACV predictions had notably higher MSE, likely due to overestimating infection extent, which was confirmed by a manual review. Foreground metrics were worse overall, with HSV and HAdV performing best, while IAV and RV performed the worst, likely due to models hallucinating the signal. U-Net generally outperformed pix2pix, except for IAV, where pix2pix produced clearer, more accurate results despite U-Net’s blurrier predictions.

## Supplementary Note 2: Cell classification metrics

Table 5 reports IoU, F1, accuracy, precision and recall metrics, but based on whole-cell masks, as opposed to previously used nuclei masks, which were available only for the HAdV and HAdV (2ch) datasets. The conclusions remain consistent: pix2pix generally outperforms U-Net, except in accuracy for both datasets and precision for the HAdV (2ch) data. Pix2pix achieves better IoU, F1, and recall scores, with precision being either comparable or slightly lower for HAdV. The accuracy, however, is lower when compared to metrics calculated from the nuclei masks. This can be attributed to the larger size of cells, which makes it easier to achieve higher IoU and recall scores. Because cells occupy a larger portion of the image, achieving high accuracy, which is heavily influenced by the background, becomes more challenging.

## Supplementary Note 3: Standard Deviation Across Samples

We calculated the standard deviations of performance metrics across all models and datasets to assess the variation in model performance. The results are detailed in Tables 6 and 7.

The analysis reveals that VACV exhibits the highest variance in both general and background metrics. This suggests that the models made substantial errors on a few samples, likely exacerbated by the smaller size of the VACV test set. In contrast, the standard deviations for background metrics across other viruses were relatively low. However, foreground metrics showed higher variability, with IAV demonstrating the greatest standard deviation in MSE.

